# Structural basis for recognition of the FLAG-tag by anti-FLAG M2

**DOI:** 10.1101/2024.03.25.586599

**Authors:** J. Wouter Beugelink, Els Sweep, Bert J.C. Janssen, Joost Snijder, Matti F. Pronker

## Abstract

The FLAG-tag/anti-FLAG system is a widely used biochemical tool for protein detection and purification. Anti-FLAG M2 is the most popular antibody against the FLAG-tag, due to its ease of use, versatility, and availability in pure form or as bead conjugate. M2 binds N-terminal, C-terminal and internal FLAG-tags and binding is calcium-independent, but the molecular basis for the FLAG-tag specificity and recognition remains unresolved.

Here we present an atomic resolution (1.17 Å) structure of the FLAG peptide in complex with the Fab of anti-FLAG M2, revealing key binding determinants. Five of the eight FLAG peptide residues form direct interactions with paratope residues. The FLAG peptide adopts a 3_10_ helix conformation in complex with the Fab. These structural insights allowed us to rationally introduce point mutations on both the peptide and antibody side. We tested these by surface plasmon resonance, leading us to propose a shorter yet equally binding version of the FLAG-tag for the M2 antibody.

## Introduction

The FLAG-tag/anti-FLAG system is a widely used biochemical tool for protein purification, (co-)immunoprecipitation (IP), Western Blot, immunocytochemistry, chromatin IP, immunohistochemistry and other (*e*.*g*. chemical biology) applications[1]. The FLAG-tag, consisting of the DYKDDDDK peptide sequence, was originally designed to be a short hydrophilic purification handle with an internal protease cleavage site (Enterokinase, also known as Enteropeptidase/TMPRSS15; - DDDDK-) for protein identification and purification[2]. The short hydrophilic nature of the FLAG peptide and the high specificity and affinity of available antibodies, combined with the possibility of gentle elution with a synthetic FLAG peptide (or, alternatively, low pH or proteolytic release), make it suitable for many applications[1,3,4].

The original anti-FLAG antibody M1 (also known as 4E11) binds calcium-dependent and only to FLAG-tags at the very N-terminus of the (mature) protein[5]. Later iterations of anti-FLAG antibodies such as M2, M5, L5 and 2H8 did not suffer from these limitations, binding calcium-independent to FLAG-tags on both N- and C-termini, or directly following a starter methionine (*e*.*g*. for cytosolic protein), as well as internal tags (*e*.*g*. embedded in a flexible linker between two domains)[5–7]. The mouse monoclonal anti-FLAG M2 in particular is widely used due to the availability of a hybridoma cell line[5], and commercial availability of purified anti-FLAG M2 as free IgG and pre-coupled affinity resins. Due to its qualities and widespread popularity, the FLAG-tag/anti-FLAG system has been the subject of intense optimization, both from the peptide and antibody side[6–12].

Previously, structures of other widely used antibody/peptide tag complexes have been described, such as for the anti-Influenza Hemaglutinin (HA)/HA-tag[13,14], anti-cMyc/cMyc-tag[15] and anti-His/His-tag[16]. Despite the wide use of the anti-FLAG-M2 antibody, neither its sequence nor the structural basis for its specific interaction with the FLAG-tag have been publicly available, hampering its application in genetically engineered affinity reagents or structure-based methods to improve binding. We recently used proteomic sequencing in combination with a previously determined incomplete high-resolution crystal structure[17] to determine the protein sequence of the anti-FLAG M2 heavy and light chains, and incorporated these sequences into mammalian expression vectors for recombinant production[18]. We validated recombinantly expressed anti-FLAG M2 using these plasmids by Western blot, yielding indistinguishable results compared to the commercially available antibody[18].

Here, we present a high-resolution structure of the anti-FLAG M2 Fab in complex with the FLAG peptide. This provides us with key residue-specific binding determinants, suggesting possible modifications. Site-directed mutagenesis of the anti-FLAG M2 and the FLAG peptide was combined with surface plasmon resonance (SPR) to test different variants of both, showing that a shorter version of the FLAG-tag is possible without impeding the affinity.

## Results

### Structure determination

To create a Fab version of our previously described recombinant anti-FLAG M2[18], the heavy chain construct was truncated in between the C_H_1 domain and the hinge region, keeping the C-terminal -Ala_3_His_8_ tag for purification purposes. This construct was co-expressed with the original light chain construct in Expi293 cells, after which the Fab was purified from the cell supernatant using Ni-affinity followed by size exclusion chromatography (see Methods for details). Crystallization screens were set up after mixing the Fab with synthetically produced FLAG peptide, yielding several crystallization hits. Our best crystal diffracted anisotropically to a maximum resolution of 1.17 Å, in the same crystal form as the original *apo* anti-FLAG M2 Fab structure[17] (table 1). Phasing by molecular replacement readily revealed strong *F*_o_-*F*_c_ difference density near the paratope region (figure 1), confirming the complex was formed in the crystal, which allowed modelling the first 6 residues of the FLAG peptide (DYKDDD, supp. video 1).

**Table 1:**
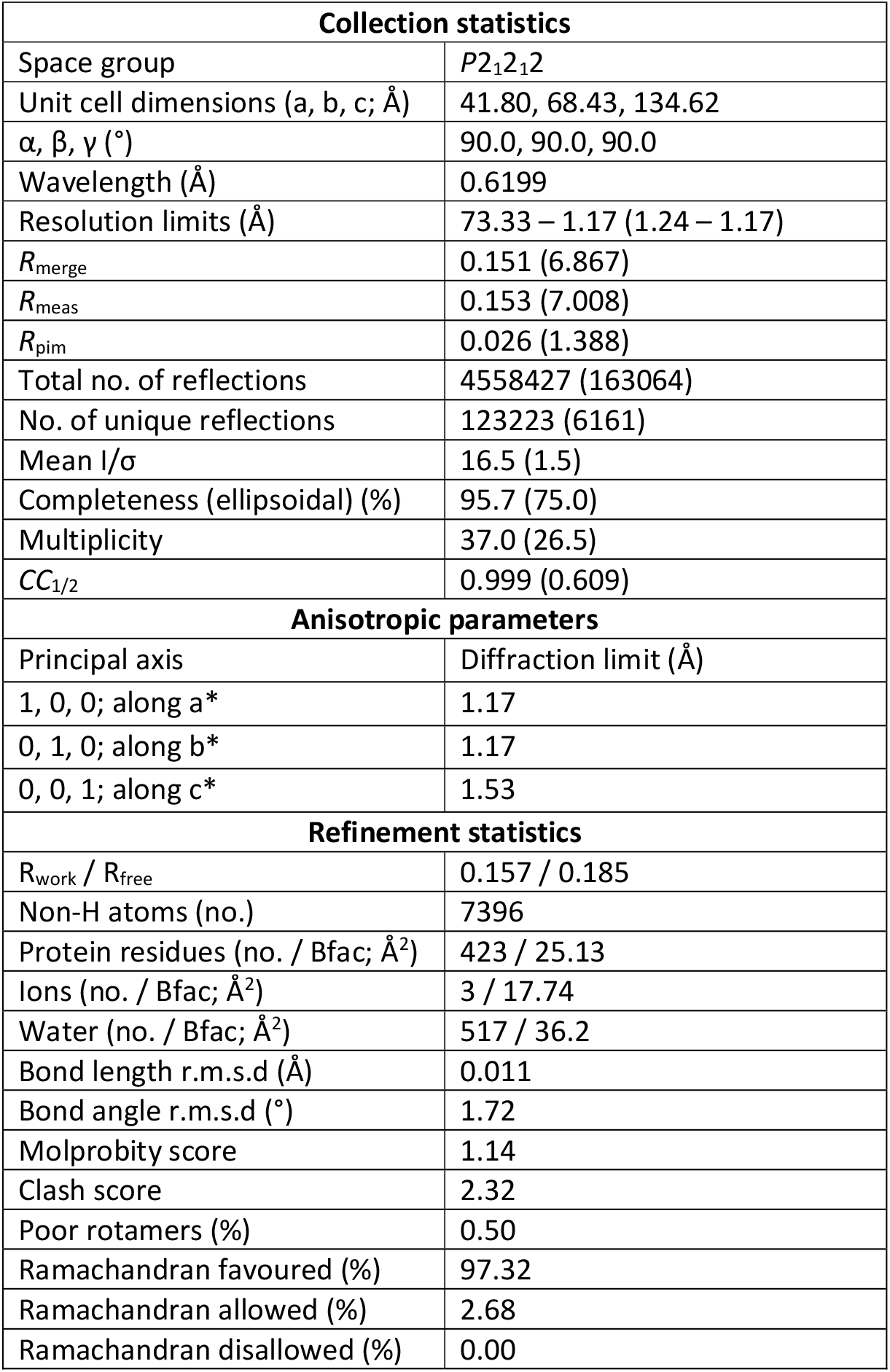
Crystallographic data collection and refinement parameters.

**Figure 1.**
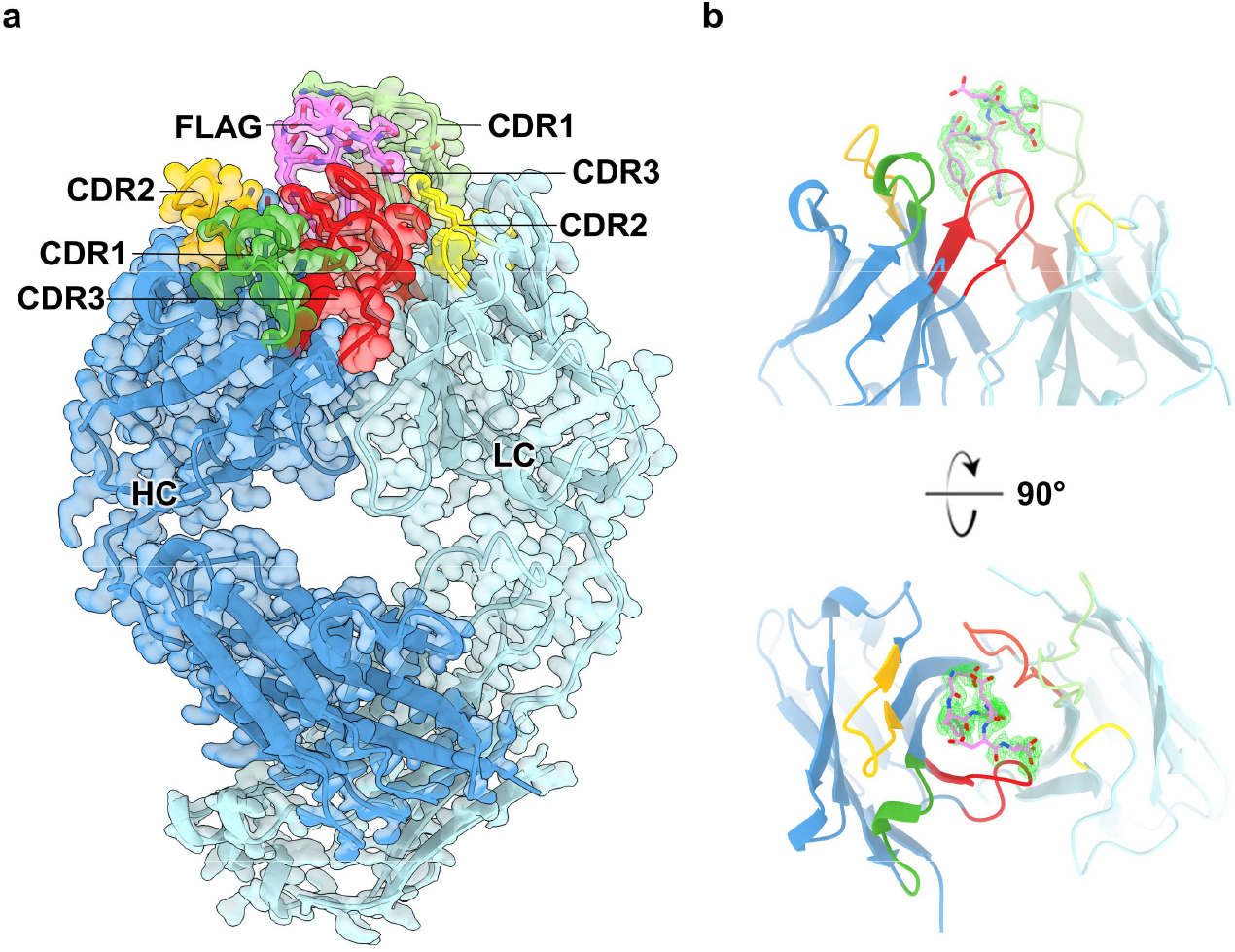
Structure of the anti-FLAG M2 Fab in complex with the FLAG peptide. A) Overview of the structure of the complex of the anti-FLAG M2 Fab and the FLAG peptide; CDR loops are indicated in green (CDR1), yellow (CDR2) and red (CDR3). (B) m*F*_o_-D*F*_c_ difference map at 3 r.m.s.d based on the (complete) *apo* anti-FLAG M2 Fab structure (PDB 7BG1) reveals the density for the FLAG peptide near the expected binding site.

### Anti-FLAG M2 binds to the FLAG-tag with both chains

As anticipated for a highly charged and hydrophilic epitope, most of the interactions formed between the FLAG peptide and anti-FLAG M2 are either through hydrogen bonds, salt bridges or ionic interactions (figure 2, table 2). Five out of the eight FLAG peptide residues (Asp1, Tyr2, Lys3, Asp4 and Asp6) appear to contribute directly to the interaction with M2. FLAG peptide residues Tyr2, Lys3 and Asp4 appear to form crucial interactions, predominantly involving M2 residues heavy chain Glu99 and light chain Arg32 (figure 2, table 2). Tyr2 and Lys3 protrude into the hole between the heavy and light chain variable domains, where they are stabilized by both hydrophilic and hydrophobic interactions. Asp1 and Asp6 form direct salt bridges with light chain paratope residues His31 and Lys55, respectively (figure 2, table 2). The backbone carbonyl of Lys3 forms a hydrogen bond with the side chain carboxamide of light chain Asn33. Similarly, the backbone carbonyl of Asp6 forms a hydrogen bond with the carboxamide side chain of light chain Asn35. FLAG residue Asp5 is oriented away from the M2 binding site and residues Asp7 and Lys8 are not resolved in the electron density, indicating that these residues do not contribute directly to interactions.

**Table 2:**
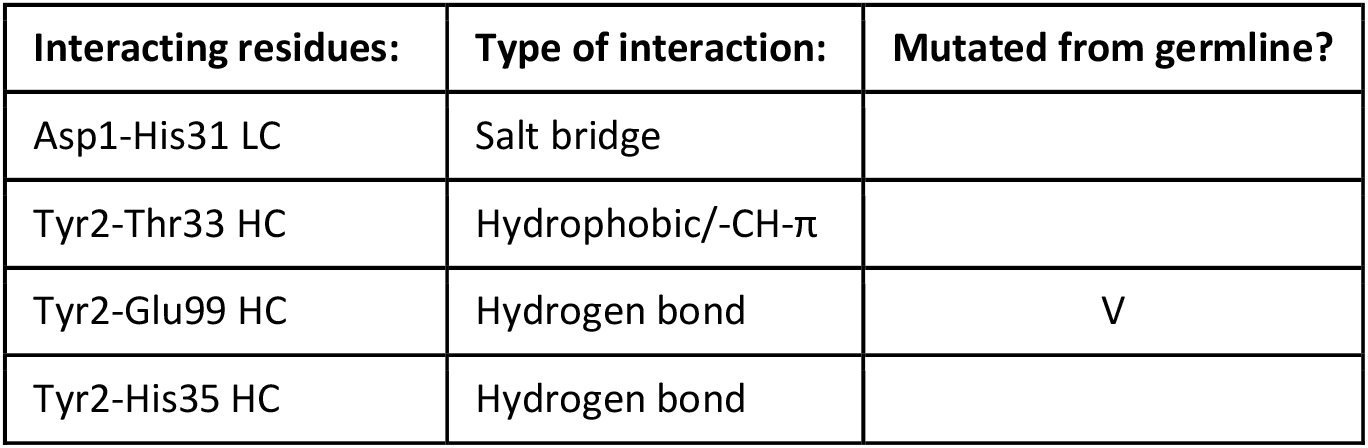
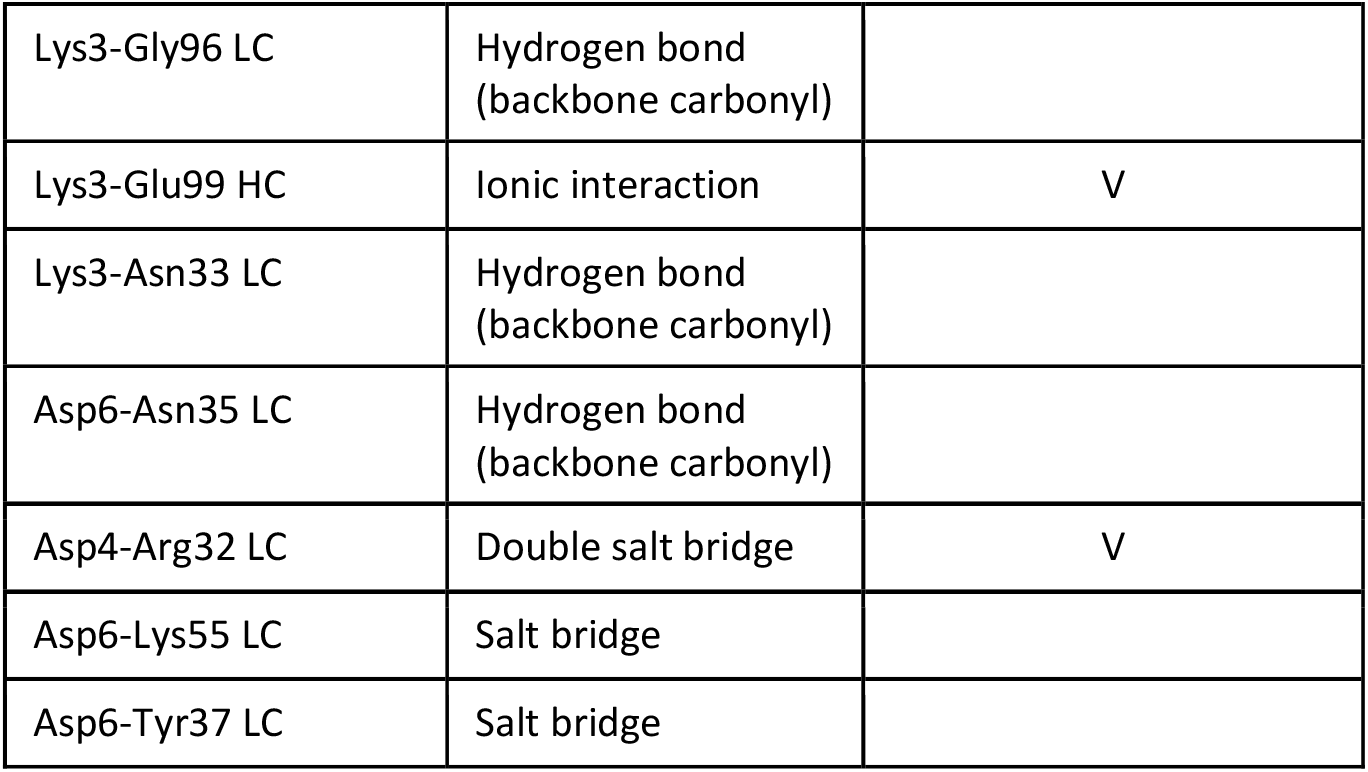
Observed interactions (LC: light chain, HC: heavy chain)

| Interacting residues: | Type of interaction: | Mutated from germline? |
| --- | --- | --- |
| Asp1-His31 LC | Salt bridge |  |
| Tyr2-Thr33 HC | Hydrophobic/-CH-π | |
| Tyr2-Glu99 HC | Hydrogen bond | V |
| Tyr2-His35 HC | Hydrogen bond |  |
| Lys3-Gly96 LC | Hydrogen bond<br>(backbone carbonyl) |  |
| Lys3-Glu99 HC | Ionic interaction | V |
| Lys3-Asn33 LC | Hydrogen bond<br>(backbone carbonyl) |  |
| Asp6-Asn35 LC | Hydrogen bond<br>(backbone carbonyl) |  |
| Asp4-Arg32 LC | Double salt bridge | V |
| Asp6-Lys55 LC | Salt bridge |  |
| Asp6-Tyr37 LC | Salt bridge |  |

**Figure 2.**
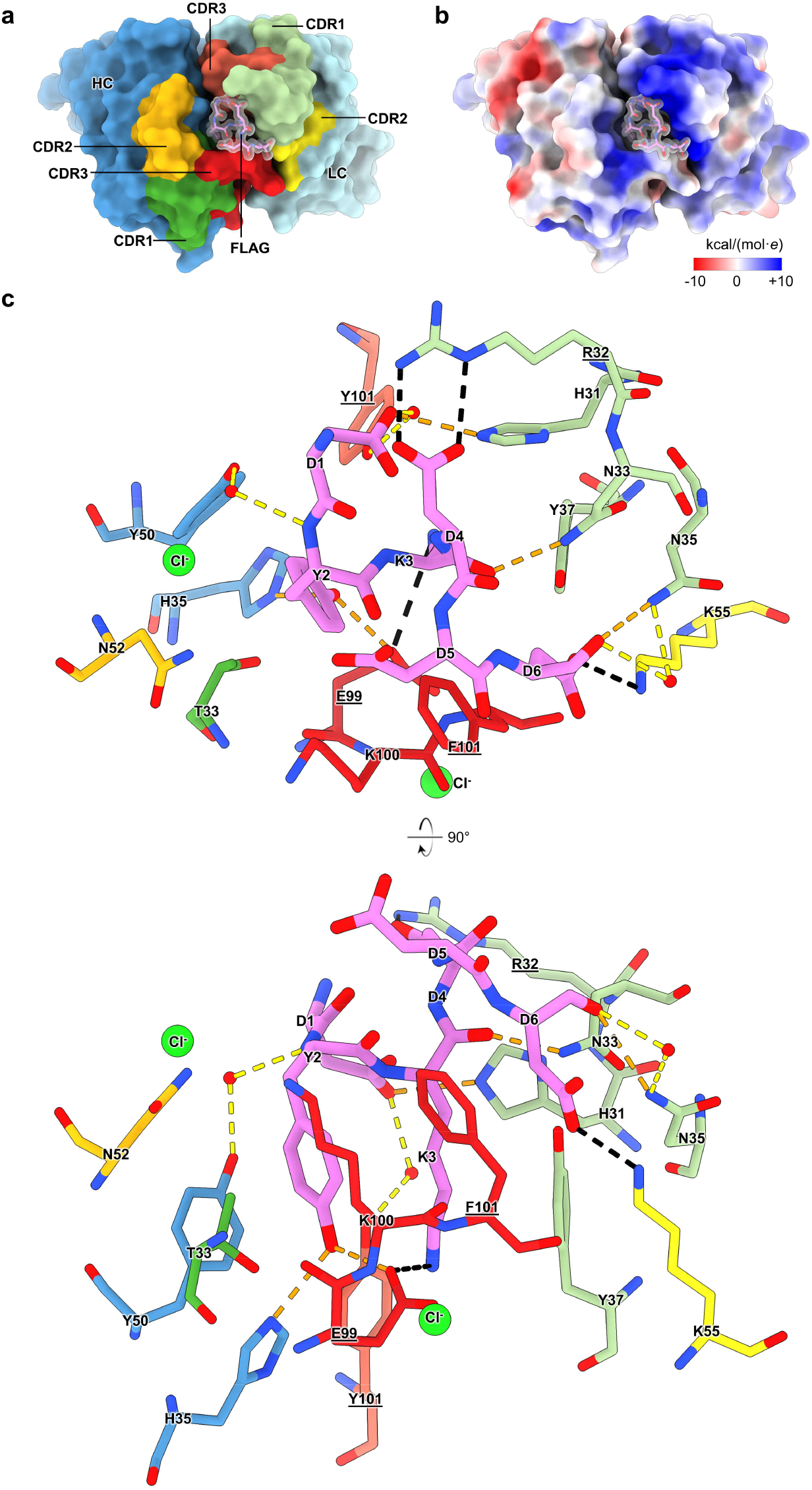
An intricate network of mostly hydrophilic interactions stabilises the interaction of the FLAG peptide with the anti-FLAG M2 Fab. A) Overview of the complex structure in surface (Fab) and sticks (FLAG peptide) representation, colouring as in figure 1 (heavy chain in dark blue, light chain in light blue, FLAG peptide in pink, CDR1 in green, CDR2 in yellow and CDR3 in red). (B) Electrostatic surface potential map of the Fab shows a mostly electropositive paratope, which would complement the net negatively charged FLAG peptide (shown as pink sticks), same orientation as in figure 2A. (C) Direct hydrogen bonds between the FLAG peptide and paratope residues are indicated as orange dashed lines, salt bridges *idem* in black, indirect hydrogen bonds mediated by a single water molecule (red spheres) are shown as yellow dashed lines. Paratope residues mutated from germline (most likely by somatic hypermutation) are underlined. Protein and peptide are coloured as in figure 1 and 2A.

Marked conformational changes in the paratope of M2 are apparent comparing the *apo* and FLAG-bound structures (figure 3, supp. video 2). Light chain Arg32 wraps around the FLAG peptide to form a double salt bridge with Asp4. Other paratope residues that substantially change their conformation upon binding are Glu99 and Phe101 of the heavy chain CDR3 loop, both of which form stabilizing interactions with the buried pair of side chains of Tyr2 and Lys3 of the FLAG sequence in the central cavity between heavy and light chains (figure 2). Interestingly, both light chain Arg32 and heavy chain Glu99 and Phe101 are all mutated compared to mouse germline ancestral sequences[19,20], likely the result of somatic hypermutation. A subtle general compaction of the N-terminal (variable) Ig domains of the heavy and light chains is also observed compared to the *apo* structure (figure 3, supp video 2).

**Figure 3.**
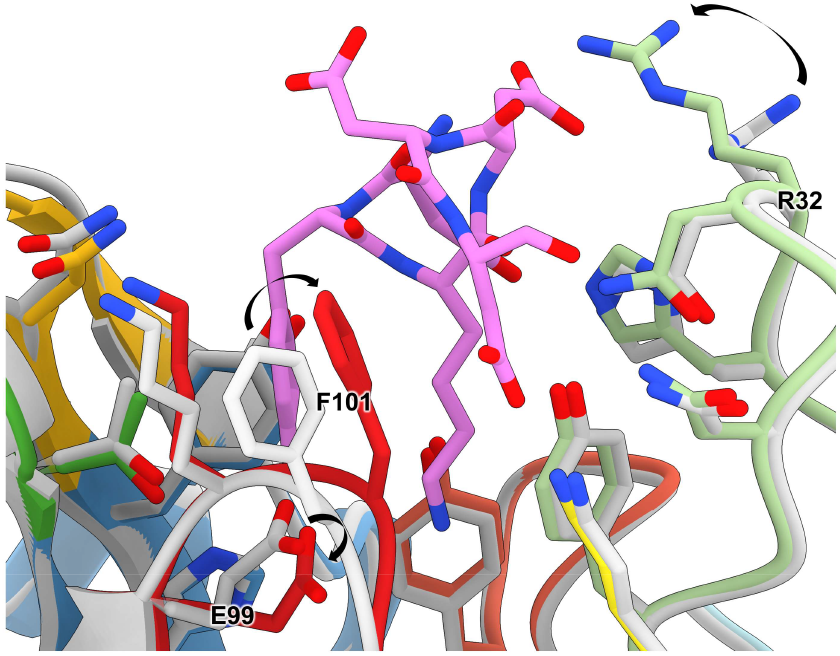
Conformational changes observed between the structure of the *apo* anti-FLAG M2 Fab and the complex with the FLAG peptide. Overlay of the *apo* (grey) anti-FLAG M2 structure (PDB 7BG1) and the complexed structure with FLAG peptide presented here (coloured as in Figure 1).

Another feature observed is that the N-terminal glutamine of the heavy chain forms a pyroglutamate, a phenomenon more commonly observed for antibodies[21,22] (supplementary figure 1). This however does not appear to affect the binding to the FLAG sequence, since the heavy chain N-terminus is distal from the FLAG peptide binding site (25 Å). Several ordered water molecules as well as two chloride ions were observed neighbouring the FLAG peptide binding site (figure 2, supplementary figure 2). Three of these water molecules simultaneously form hydrogen bonds with both paratope and FLAG peptide, mediating indirect interactions between residues FLAG Asp1 and light chain Tyr101, FLAG Tyr2 and heavy chain Tyr50 and between FLAG Asp6 and light chain Asn35.

### The FLAG peptide adopts a 3_10_ helix conformation

Unexpectedly, we observed that the FLAG peptide adopts a 3_10_ helix conformation with two full turns (figure 4c). Whereas in α-helices backbone hydrogen bonds are formed between residues *i* and *i*+4, we see the typical tighter wound 3_10_ helix backbone hydrogen bonding interactions of residue *i* and *i*+3 for the pairs of Asp1-Asp4, Tyr2-Asp5 and Lys3-Asp6 in the FLAG peptide. While a 3_10_ helix is typically energetically less favourable than a regular α-helix, it appears that the FLAG sequence was inadvertently designed with a preference for 3_10_ helix formation[23], with aspartate residues in position 1 and 4. Indeed, both these aspartates form stabilising interactions with their carboxylate side chains to the backbone nitrogen atoms of the N-terminus (Asp4) of the peptide and the backbone amide of Lys3 (Asp1). This suggests that, while not essential for anti-FLAG M2, having the FLAG-tag at the very N-terminus of the protein (without an initiator methionine) might still be beneficial in lowering the energetic penalty for forming a 3_10_ helix, compared to an internal or C-terminal FLAG-tag. Since the N-terminal amino group carries a formal charge at neutral pH and is expected to form a stronger interaction with the carboxylic acid of the side chain of Asp4 compared to an amide nitrogen for an internal or C-terminal tag, it could favour the 3_10_ helix conformation observed in our complex structure.

**Figure 4.**
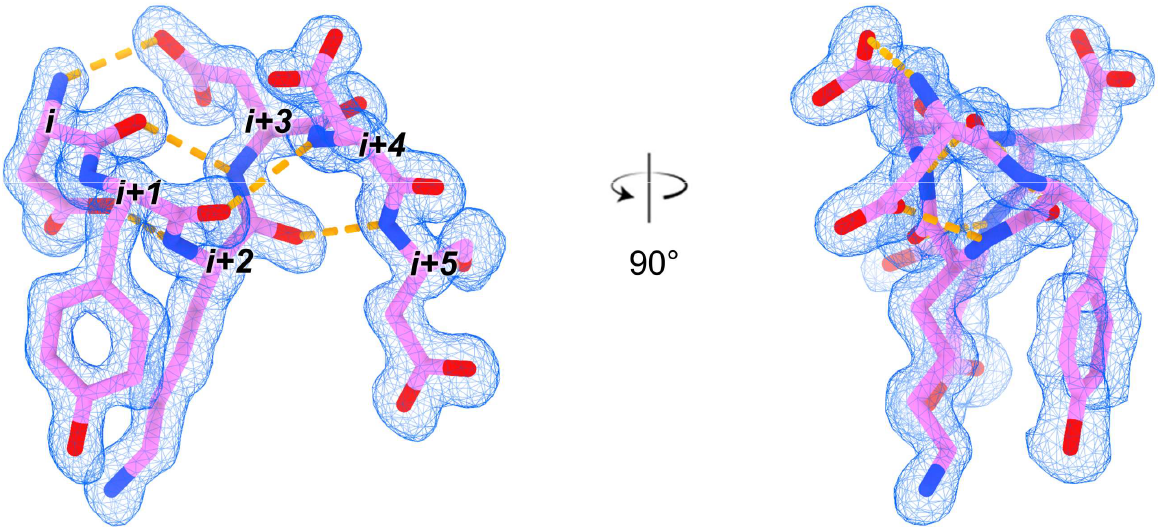
The FLAG peptide adopts a 3_10_ helix conformation in complex with anti-FLAG M2. Backbone hydrogen bonds within the FLAG peptide are indicated as orange dashed lines. 2m*F*_o_-D*F*_c_ electron density at 1.3 r.m.s.d. shown as blue mesh.

### Introducing an extra salt bridge between FLAG-tag Asp5 and heavy chain Lys100

#### Surface plasmon resonance to determine the affinity of wildtype and variant FLAG sequences

Since the side chain of Asp5 is not involved in direct interactions with the antibody paratope, we considered this position to be amenable to mutation with the goal of enhancing the binding affinity. The side chain of heavy chain Lys100 in CDR3 is close to the Asp5 side chain carboxylic acid, but still too far for a direct salt bridge in our structure (6.5 Å, figure 5a). We reasoned that mutating either FLAG-tag Asp5 to glutamate or heavy chain Lys100 to arginine, or combining those two mutations would bring their side chains in close enough proximity in the complex to form an extra stabilizing salt bridge, thereby enhancing the binding affinity.

**Figure 5.**
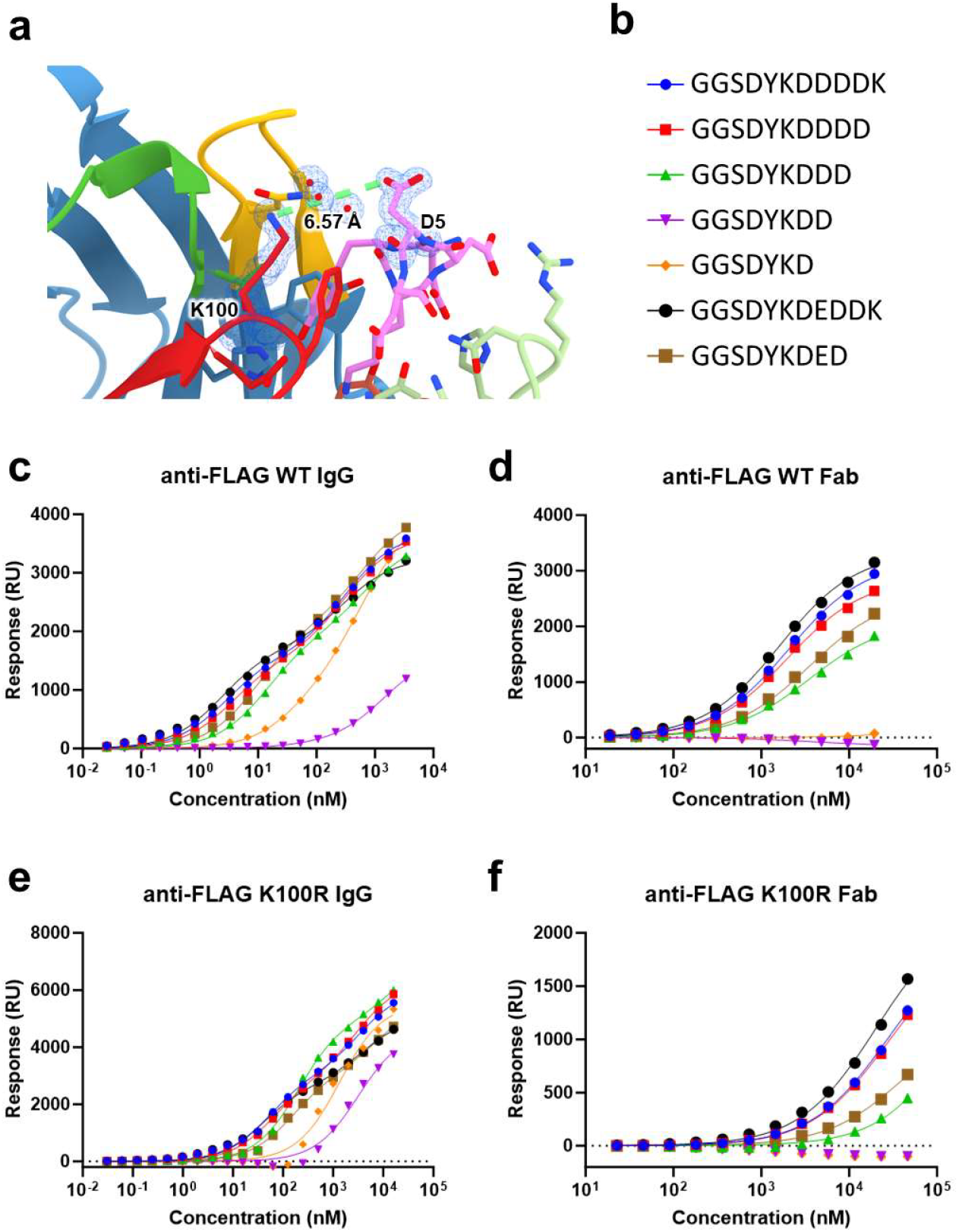
Surface plasmon resonance quantifies binding affinities of the FLAG/anti-FLAG M2 interaction and variants. A) Location of Fab heavy chain K100 and FLAG peptide D5, displaying a distance of 6.6 Å. Selected part of 2m*F*_o_-D*F*_c_ electron density at 1.3 r.m.s.d. shown as blue mesh. (B) Sequences of FLAG peptides used for affinity measurements. Coloured shapes correspond to curves in c-f. (C) Binding of anti-FLAG WT IgG to FLAG peptides. (D) Binding of anti-FLAG WT Fab to FLAG peptides. (E) Binding of anti-FLAG K100R IgG to FLAG peptides. (F) Binding of anti-FLAG K100R Fab to FLAG peptides.

A series of N-terminally biotinylated synthetic FLAG peptide variants were immobilised to streptavidin-coated SPR chips to determine the binding affinity of anti-FLAG M2. C-terminally truncated variants, as well as two variants with a glutamate in place of Asp5 of the FLAG sequence were tested (DYKDEDDK and DYKDED, table 3). Anti-FLAG M2 Fab and full IgG were used in the mobile phase; the Fab was used to obtain 1:1 binding following a Langmuir isotherm model similar to purification and (co-)IP applications (for monomeric target proteins), the full IgG to quantify avidity-enhanced binding more akin to usage of the antibody in Western blot, immunocytochemistry and immunohistochemistry.

**Table 3:** Surface plasmon resonance data.

|  | Anti-FLAG WT IgG |  | Anti-FLAG<br>WT Fab | Anti-FLAG K100R IgG |  | Anti-FLAG<br>K100R Fab |
| --- | --- | --- | --- | --- | --- | --- |
| | High<br>affinity<br>(nM) | Low<br>affinity<br>(nM) | One site<br>(μM) | High affinity<br>(nM) | Low<br>affinity<br>(μM) | One site<br>(μM) |
| GGSDYKDDDDK | $2.95 \pm 0.31$ | $306 \pm 8.9$ | $1.91 \pm 0.24$ | $59.8 \pm 7.1$ | $4.30 \pm 0.52$ | $23.9 \pm 3.0$ |
| GGSDYKDDDD | $3.08 \pm 0.40$ | $289 \pm 14$ | $2.01 \pm 0.43$ | $61.3 \pm 1.8$ | $4.04 \pm 6.0$ | $25.3 \pm 3.0$ |
| GGSDYKDDD | $9.72 \pm 2.6$ | $512 \pm 130$ | $3.52 \pm 0.72$ | $197 \pm 13$ | $16.7 \pm 2.8$ | $118 \pm 73$ |
| GGSDYKDD | $785 \pm 190$ | - | n.b. | $5250 \pm 1300$ | - | n.b. |
| GGSDYKD | $288 \pm 48$ | - | n.b. | $1460 \pm 220$ | - | n.b. |
| GGSDYKDEDDK | $2.23 \pm 0.25$ | $274 \pm 30$ | $1.78 \pm 0.13$ | $45.9 \pm 4.8$ | $4.08 \pm 0.20$ | $18.3 \pm 2.1$ |
| GGSDYKDED | $10.2 \pm 0.48$ | $555 \pm 32$ | $3.54 \pm 0.24$ | $131 \pm 15$ | $6.99 \pm 1.1$ | $62.1 \pm 29$ |

As expected, the apparent affinities were orders of magnitude higher for the full IgG (*K*_*D*_ of 2.95 ± 0.31 nM and 306 ± 8.9 nM for the wildtype peptide, table 3, figure 5, supplementary figures 3 and 4), where a two-site binding model was used to fit the data, compared to the Fab (*K*_*D*_ of 1.91 ± 0.24 μM, one-site binding model). However, the trends between the preference for different FLAG-tag variants were highly similar between the Fab and full IgG data. Truncating the original FLAG sequence DYKDDDDK with one residue from the C-terminal side did not affect the binding affinity, but shorter truncations (in particular the extra short 4- and 5-residue variants DYKD and DYKDD) did dramatically suffer in their binding affinity (52.9 ± 22.1 nM and 154 ± 100 nM, respectively).

The Asp5Glu mutation did appear to subtly benefit binding for the full wt anti-FLAG M2 IgG, with a *K*_*D*_ of 2.95 ± 0.31 nM and 2.23 ± 0.25 nM for the wt and mutated peptide, respectively (p=0.01, table 3 and figure 5). However, no significant difference was observed when comparing a truncated peptide, which measured 9.72 ± 2.6 nM and 10.2 ± 0.48 nM for the wt and mutant peptide, nor did the wt anti-FLAG M2 Fab show a significant difference in binding affinity between wt and Asp5Glu FLAG peptide (1.91 ± 0.24 μM and 1.78 ± 0.13 μM, respectively; p=0.19).

When using the heavy chain Lys100Arg mutated Fab or IgG in the mobile phase, the affinities dropped by about 20-fold for both the original FLAG sequence and the other variants we tested (table 3, figure 5, supplementary figure 3 and 4). Possibly, the Lys100Arg mutation locally changes the fold and/or surface properties of the antibody in a way that is disadvantageous for binding the FLAG peptide (either wildtype or Asp5Glu mutant).

## Discussion

Our high-resolution structure of the FLAG/anti-FLAG M2 Fab complex reveals the structural determinants of the interaction at the basis of this widely used biochemical tool. It also represents the highest resolution structure of an antibody-antigen complex in the protein databank to date.

We only observed well-resolved electron density for the first six out of eight residues of the FLAG peptide (density observed for DYKDDD compared to the full sequence DYKDDDDK), prompting us to determine binding affinities for shorter truncated versions of the FLAG sequence. While truncating the FLAG peptide by deleting the C-terminal lysine residue did not appear to impair binding, shorter truncations did suffer in terms of binding affinity. This is particularly surprising for the six-residue truncated version (DYKDDD), since no density is observed for Asp7, suggesting it does not directly contribute to the binding affinity. Possibly, having the C-terminus at position six (DYKDD<u>D</u>DK) interferes with 3_10_ helix formation or complex formation by having an extra negative charge so close to the epitope/paratope interaction. Of note, while a shorter (*e*.*g*. 6- or 7-residue) C-terminally truncated version of the FLAG-tag might be beneficial for certain applications, one has to bear in mind that by truncating any C-terminal residue from the original FLAG-tag sequence, the enterokinase/enteropeptidase/TMPRSS15 proteolytic cleavage motif (DDDDK) will be lost.

Now that we determined the structural basis of the FLAG/anti-FLAG M2 interaction, it is interesting to revisit previous empirical attempts to determine the binding determinants of this interaction[9,10]. Indeed, these studies also found that residues 5, 7 and 8 of the FLAG sequence appear to contribute little to the affinity for anti-FLAG M2. Substituting Asp4 and Asp6 with glutamate still permitted binding[10], which could be explained by these residues still having a carboxylic acid side chain which can still form similar salt bridges to the original aspartate residues, if some flexibility in the peptide or paratope would permit the accommodation of the additional aliphatic -C_γ_H_2_-. According to a high-throughput phage display screen[9], the main binding determinants in the FLAG sequence are Tyr2, Lys3 and Asp6, similar to what we conclude based on the structure. Asp4 and, in particular, Asp1 were also enriched, whereas position 5, 7 and 8 were completely random, in agreement with their lack of contribution to the binding interface in our structure.

In conclusion, our data provide a structural framework for understanding the interaction of the FLAG-tag with its most used antibody, anti-FLAG M2, in agreement with previous empirical studies. With the anti-FLAG M2 sequence now publicly available and its interactions with the FLAG-tag defined with atomic detail, the stage is set for further structure-based optimisation of FLAG-tag based affinity reagents.

## Materials and Methods

### Construct design

The heavy chain Fab construct for anti-FLAG M2 was generated by deletion PCR and in vivo assembly (IVA) cloning[24], truncating after Gly219 using the following primers: gcggccgcaCATCACCACCATCATCACCATCATTGATAAC (forward) and GTGATGTGCGGCCGCGCCACAGTCGCGCGGCAC (reverse), to ensure an in-frame NotI site (translates to three alanines with an extra adenosine base) followed by the octahistidine tag for purification, both already present in the original full IgG anti-FLAG M2 heavy chain plasmid[18].

The K100R mutation was introduced into the heavy chain by site-directed mutagenesis using IVA cloning, using the following primers: CTATTGTGCGCGAGAGagaTTCTATGGTTACGATTATTGGGGCCAAG (forward) and tctCTCTCGCGCACAATAGTAGACAGCACTATCC (reverse). The Fab of this mutant heavy chain construct was generated in the same way as the wildtype version (see above).

### Protein expression

The IgG and Fab constructs were transiently expressed by mixing two plasmids encoding for the heavy and light chain in a 1:1 (m/m) ratio and transfecting this mixture into Expi293 cells. To this end, plasmid mixture at 30 μg/mL and Polyethylenimine “Max” (PEI MAX®) at 90 μg/mL were combined with the Expi293 cells at 3*10^6^ viable cells/mL in a 1:1:28 (v/v/v) ratio. First the plasmid mixture was dropwise added to the PEI MAX®, and after 30 minutes of incubation this new mixture was dropwise added to the cells. The final concentrations of plasmid and PEI MAX® were thus 1 and 3 μg/mL, respectively. After 6 days the proteins were harvested by spinning down the culture twice for 10 min at 4000 rpm and subsequently filtering the medium using a 0.22 μm filter.

### Protein purification

Full IgG and Fab versions of anti-FLAG M2 were purified from the Expi293 cell supernatant by a combination of immobilized metal affinity chromatography (IMAC) followed by size exclusion chromatography (SEC). Filtered cell supernatant was loaded onto a 5 mL HisTrap-excel column (Cytiva) equilibrated with IMAC A buffer (500 mM NaCl, 25 mM HEPES pH 7.8) at a flowrate of 5 ml/min. The column was washed with 200-300 mL of either 3% (v/v, Fab) or 5% (v/v, IgG) IMAC B (500 mM NaCl, 500 mM imidazole, 25 mM HEPES pH 7.8) in IMAC A, until the UV A280 signal reached a stable baseline. Protein was eluted with 40% IMAC B in IMAC A (wildtype Fab) or 100% IMAC B (K100R Fab and both IgG’s). The IMAC eluates were concentrated to 1-1.5 ml using 10 kDa MWCO concentrators (Amicon), before injection onto a HiLoad 16/600 superdex200 SEC column (Cytiva) equilibrated with SEC buffer (150 mM NaCl, 20 mM HEPES pH 7.5). Fab and IgG peak fractions were pooled and concentrated with 10 kDa MWCO concentrators (Amicon) to a concentration of 2-6 mg/ml.

### Crystallization

Fab-FLAG peptide complexes were mixed in a 1:3.5 molar ratio at a protein concentration of 5.6 mg/ml. This was used to set up sitting drop vapour diffusion crystallization screens, mixing 150 nL protein/peptide complex sample with 150 nL reservoir solution, at 293 K. Our best diffracting crystal grew in a condition of 0.1 M Tris-HCl pH 8, 20% (w/v) polyethylene glycol 6000, 0.2 M NH_4_Cl. Crystals were cryo-protected with reservoir solution supplemented with 25% (v/v) glycerol before plunge-freezing in liquid nitrogen.

### Data collection and structure determination

Data was collected at Diamond Light Source beamline I24, equipped with a CdTe Eiger2 9M detector, at a wavelength of 0.6199 Å. Due to the anisotropic nature of the data, three datasets collected on the same crystal were integrated and merged using the multi-autoPROC+STARANISO[25] pipeline, as integrated in Diamond ISPyB[26]. Data was then imported into CCP4i2[27]. Given that the crystallization condition and determined unit cell closely resembled that of the *apo* structure (PDB 2G60[17]/7BG1[18]), FreeR flags were copied from this dataset and extended to the higher attained resolution. The *apo* anti-FLAG M2 Fab structure (PDB 7BG1)[18] was used for molecular replacement using PHASER[28]. Density for the FLAG-tag peptide was observed, and the peptide was manually built, after adjusting CDR loops where necessary, in COOT[29]. The structure was then refined using iterative rounds of manual adjustment in COOT and refinement in REFMAC5[30], with the setting “VDWR” set to 2.0. Quality of the geometry was analysed using MolProbity[31]. All programs were used as implemented in CCP4i2 v1.1.0[27].

### Surface plasmon resonance

N-terminally biotinylated synthetic FLAG peptide variants were ordered from Genscript. These were dissolved in SPR buffer (150 mM NaCl, 20 mM HEPES pH 7.4, 0.005% (v/v) Tween20) by rotating overnight at room temperature, followed by pH adjustment with NaOH to pH 7-8 and sonication. Peptides were printed on a streptavidin-coated SPR chip (P-Strep for full IgG, G-Strep for Fab, Sens B.V.) using a continuous flow microfluidics spotter (Wasatch) and flowing for 1 hour at RT, after which the chips were washed with SPR buffer for 10 minutes and subsequently quenched with 10 mM biotin in SPR buffer. SPR experiments were performed using an IBIS-MX96 system (IBIS technologies) at 298 K with SPR buffer as the running buffer. Chips were always washed overnight with SPR buffer to have a stable baseline. Analytes were then injected in 2× dilution series in SPR buffer, measuring from low to high concentrations, at a constant temperature of 298 K. Data were analysed using *SPRINTX* (IBIS technologies), extracted in Scrubber2 (Biologic Software), and fit to a one-site or two-site specific binding model in GraphPad Prism (Dotmatics). In short, background signal from adjacent regions without ligand bound were subtracted from signal in regions of interest, and signal was set to zero at the start of the injection series. Injections were then overlayed, and the average signals after reaching equilibrium were plotted against the analyte concentration and fitted with relevant Langmuir models.

## Acknowledgements

Diffraction experiments were performed at beamline I24 at the Diamond Light Source (DLS) synchrotron, Harwell, United Kingdom. We are grateful to local contacts at DLS for assistance in using beamline I24. This research was funded by the Dutch Research Council NWO Gravitation 2013 BOO, Institute for Chemical Immunology (ICI; 024.002.009) to J.S and NWO grant OCENW.KLEIN.026 to B.J.C.J.

## Data availability

The coordinates and structure factors of the crystal structure of the FLAG/anti-FLAG M2 complex have been deposited at the protein databank under accession code 8RMO.

## CRediT authorship contribution statement

**J. Wouter Beugelink:** Investigation, Methodology, Formal analysis, Data curation, Validation, Visualization, Project administration, Writing – review & editing. **Els Sweep:** Investigation, Methodology. Writing – review & editing. **Bert J.C. Janssen:** Formal analysis, Validation, Resources, Supervision, Writing – review & editing, Funding acquisition. **Joost Snijder:** Resources, Supervision, Writing – review & editing, Funding acquisition. **Matti F. Pronker:** Conceptualization, Investigation, Methodology, Formal analysis, Data curation, Validation, Visualization, Project administration, Writing – original draft, Writing – review & editing.

## Supplementary figure legends

Supplementary figure 1: The N-terminus of the Fab heavy chain has a pyroglutamate modification.

2m*F*_o_-D*F*_c_ electron density at 2.1 r.m.s.d. shown as blue mesh.

Supplementary figure 2: Two chloride ions and several water molecules are proximal to the FLAG peptide binding site.

Hydrogen bonds formed by water molecules are indicated as blue dashed lines. Water and chloride 2m*F*_o_-D*F*_c_ electron density at 0.9 r.m.s.d. shown as blue mesh.

Supplementary figure 3: Surface plasmon resonance sensograms show binding of antibodies to peptides.

Response over time after injecting a concentration range of anti-FLAG WT IgG, anti-FLAG WT Fab, anti-FLAG K100R IgG, and anti-FLAG K100R Fab as measured in a region of interest containing peptide with the sequence (A) GGSDYDDDDK, (B) GGSDYDDDD, (C) GGSDYDDD, (D) GGSDYDD, (E) GGSDYD, (F) GGSDYDEDDK, or (G) GGSDYDED. One replicate with a nominal spotting concentration of 200 nM is shown per experiment. Values that were averaged and used for making response curves are indicated with a red block.

Supplementary figure 4: Surface plasmon resonance response curves and fitting parameters used to quantify binding affinities of the FLAG/anti-FLAG M2 interaction and variants.

Response curves for four technical replicates at nominal spotting concentrations of 500, 200, 50, and 20 nM, for injecting a concentration range of anti-FLAG WT IgG, anti-FLAG WT Fab, anti-FLAG K100R IgG, and anti-FLAG K100R Fab as measured in a region of interest containing peptide with the sequence (A) GGSDYDDDDK, (B) GGSDYDDDD, (C) GGSDYDDD, (D) GGSDYDD, (E) GGSDYD, (F) GGSDYDEDDK, or (G) GGSDYDED. Inset tables for each graph show *K*_*D*_ and Bmax modelled for one or two binding modes per curve.

## Supplementary video legends

Supplementary video 1:

Density and model for the structure of the complex of the anti-FLAG M2 Fab with the FLAG peptide

Supplementary video 2:

Conformational changes observed in the anti-FLAG M2 paratope upon binding of the FLAG peptide.

