## Supplementary figures for "Structural basis for recognition of the FLAG-tag by anti-FLAG M2"

Supplementary figure 1: The N-terminus of the Fab heavy chain has a pyroglutamate modification.

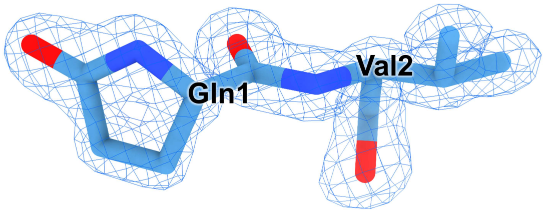


2m$F_o$-D$F_c$ electron density at 2.1 r.m.s.d. shown as blue mesh.

Supplementary figure 2: Two chloride ions and several water molecules are proximal to the FLAG peptide binding site.

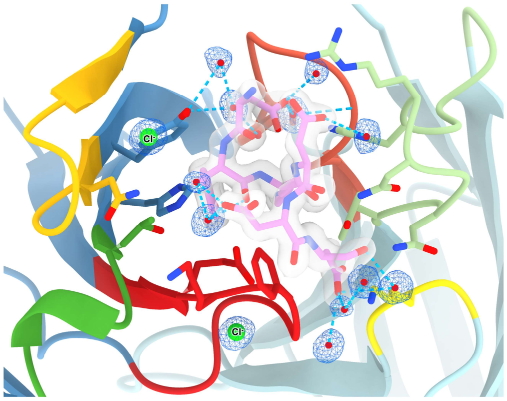


Hydrogen bonds formed by water molecules are indicated as blue dashed lines. Water and chloride 2m$F_o$-D$F_c$ electron density at 0.9 r.m.s.d. shown as blue mesh.

Supplementary figure 3: Surface plasmon resonance sensograms show binding of antibodies to peptides.

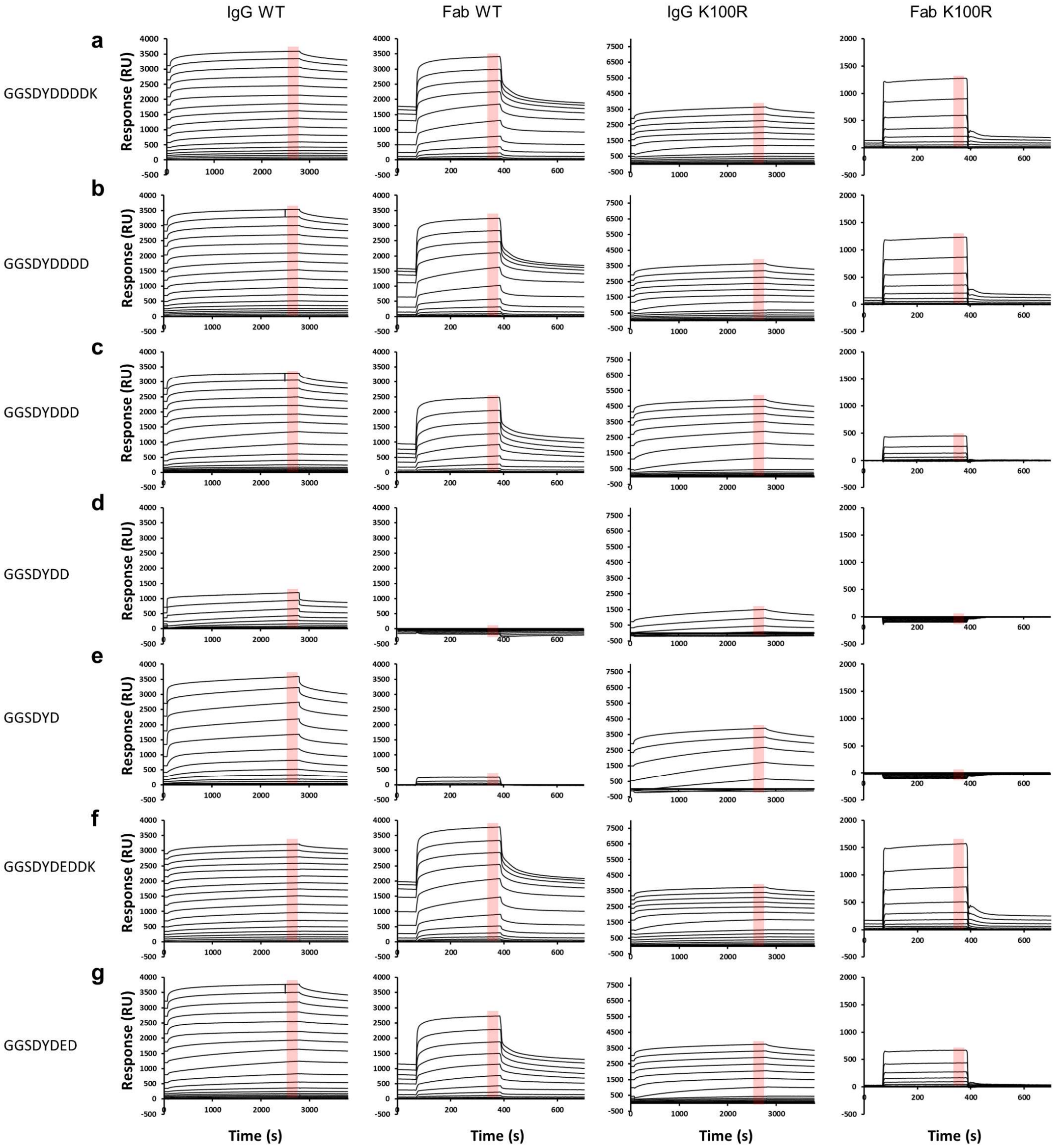


Response over time after injecting a concentration range of anti-FLAG WT IgG, anti-FLAG WT Fab, anti-FLAG K100R IgG, and anti-FLAG K100R Fab as measured in a region of interest containing peptide with the sequence (A) GGSDYDDDDK, (B) GGSDYDDDD, (C) GGSDYDDD, (D) GGSDYDD, (E) GGSDYD, (F) GGSDYDEDDK, or (G) GGSDYDED. One replicate with a nominal spotting concentration of 200 nM is shown per experiment. Values that were averaged and used for making response curves are indicated with a red block.

Supplementary figure 4: Surface plasmon resonance response curves and fitting parameters used to quantify binding affinities of the FLAG/anti-FLAG M2 interaction and variants.

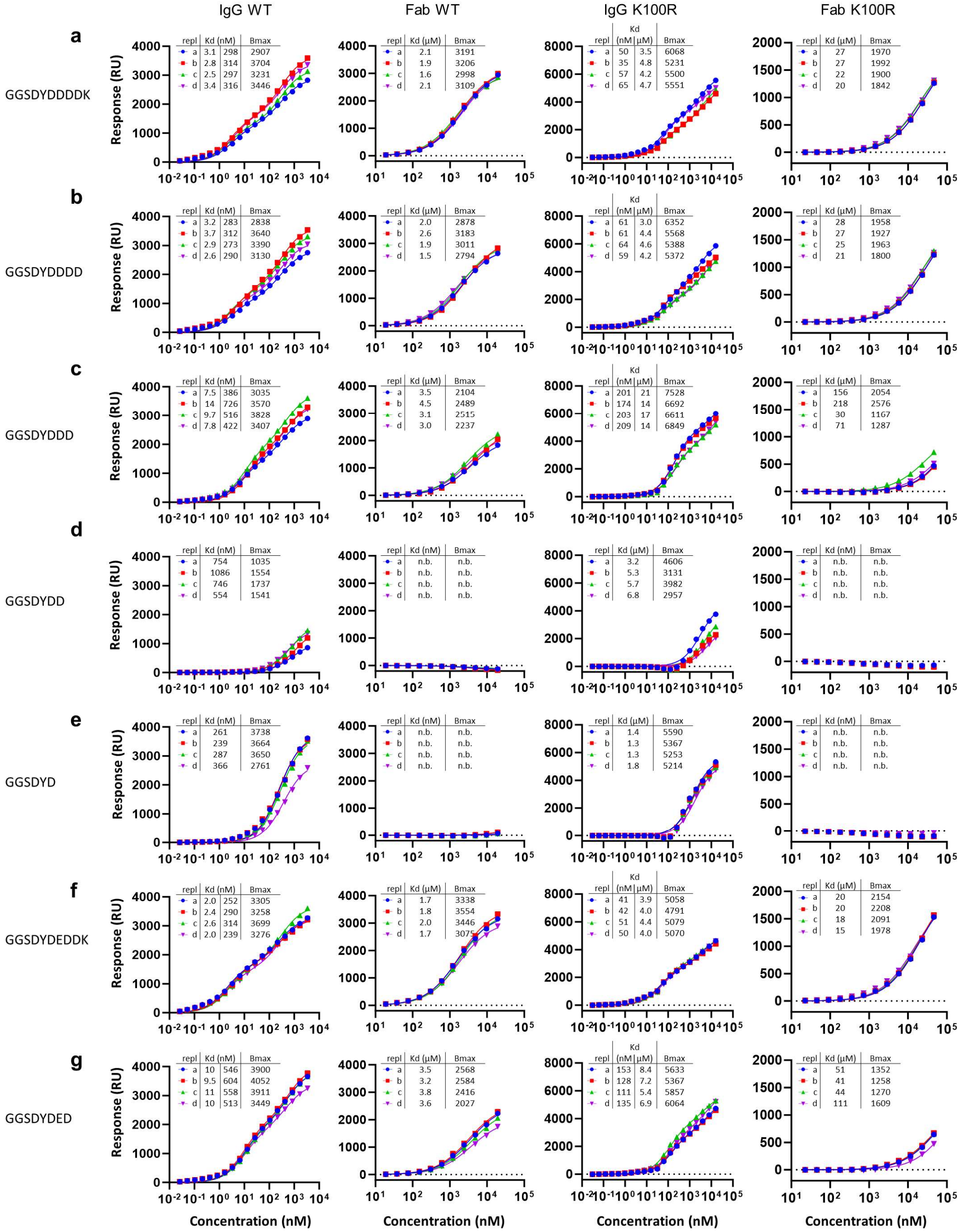

Response curves for four technical replicates at nominal spotting concentrations of 500, 200, 50, and 20 nM, for injecting a concentration range of anti-FLAG WT IgG, anti-FLAG WT Fab, anti-FLAG K100R IgG, and anti-FLAG K100R Fab as measured in a region of interest containing peptide with the sequence (A) GGSDYDDDDK, (B) GGSDYDDDD, (C) GGSDYDDD, (D) GGSDYDD, (E) GGSDYD, (F) GGSDYDEDDK, or (G) GGSDYDED. Inset tables for each graph show $K_D$ and Bmax modelled for one or two binding modes per curve.
